# Rescue of neurogenesis and age-associated cognitive decline in SAMP8 mouse: role of transforming growth factor alpha

**DOI:** 10.1101/2023.01.14.524036

**Authors:** Ricardo Gómez-Oliva, Sergio Martínez-Ortega, Isabel Atienza-Navarro, Samuel Domínguez-García, Carlos Bernal, Noelia Geribaldi-Doldán, Cristina Verástegui, Abdellah Ezzanad, Rosario Hernández-Galán, Pedro Núnez-Abades, Monica Garcia-Alloza, Carmen Castro

## Abstract

Neuropathological aging is associated with memory impairment and cognitive decline, and affects several brain areas including the neurogenic niche of the dentate gyrus of the hippocampus (DG). In the healthy brain homeostatic mechanisms regulate neurogenesis in the DG to facilitate the continuous generation of neurons from neural stem cells (NSC). Nevertheless, aging reduces the number of activated neural stem cells, and diminishes the number of newly generated neurons. Strategies that promote neurogenesis in the DG may improve cognitive performance in the elderly resulting in the development of treatments to prevent the progression of neurological disorders in the aged population.

Our work is aimed to discover targeting molecules to be used in the design of pharmacological agents to prevent the neurological effects of pathological aging. We study the effect of age on hippocampal neurogenesis using the SAMP8 mouse as a model of pathological aging. Thus, we show that in six-month-old SAMP8 mice, episodic and spatial memory are impaired, concomitantly the generation of neuroblasts and neurons is reduced and the generation of astrocytes is increased in this model. The novelty of our work resides in the fact that treatment of SAMP8 mice with a TGF-alpha targeting molecule, prevents the observed defects, positively regulating neurogenesis and improving cognitive performance. This compound facilitates the release of TGF-alpha in vitro and in vivo and activates signaling pathways initiated by this growth factor. We conclude that targeting the release of TGF-alpha may be the basis of pharmacological drugs to counteract the neurological effects of pathological aging.

## INTRODUCTION

Increasing age constitutes a risk factor for developing neurodegenerative disorders (Murman, 2015). In some individuals, age differentially affects distinct brain regions, mainly influencing memory (Gray & Barnes, 2015). Concretely, episodic memory is one of the most affected in the aging population (Nyberg, Lovden, Riklund, Lindenberger, & Backman, 2012). A form of plasticity that may help protect the brain during the continuous process of aging is the ability of some structures, including the hippocampal dentate gyrus (DG), to generate new neurons. Neurogenesis -the generation of neurons from neural stem cells-has been observed in the adult brain of many mammalian species, including humans, and throughout the individual’s life (Aimone et al., 2014). The generated neurons influence cognitive ability by affecting tasks such as memory, learning and pattern separation (Aimone, Deng, & Gage, 2011; Deng, Aimone, & Gage, 2010). Although there still exists some controversy regarding the existence of neurogenesis in the adult human hippocampus and about its role in pathological aging (Moreno-Jimenez et al., 2019; Sorrells et al., 2018), the development of new protocols (Flor-Garcia et al., 2020) have led to additional studies that describe the existence of neurogenesis throughout adult life (Moreno-Jimenez et al., 2019; Tobin et al., 2019) and indicate that this capacity is altered in patients with age-associated neurodegenerative disorders (Marquez-Valadez, Rabano, & Llorens-Martin, 2022; Terreros-Roncal et al., 2021). The generation of neurons from neural stem cells (NSC) requires an adequate physiological environment built up of cells, trophic factors, elements of the extracellular matrix, etc. that activate NSC and direct their fate towards neurons. This environment constitutes what is called a neurogenic niche. Like the rest of the organism, neurogenic niches age altering and reducing the ability of NSCs to generate neurons (Diaz-Moreno et al., 2018; Encinas et al., 2011). In general, several pieces of evidence show that hippocampal-dependent cognitive abilities decline with age in mammals, including humans, with a concomitant reduction in adult hippocampal neurogenesis (Akers et al., 2014; Magavi, Leavitt, & Macklis, 2000; Yassa et al., 2011). A key point in the regulation of neurogenesis within neurogenic niches is the control of NSC activation. NSCs may enter a prolonged quiescent state or enter an active state. NSCs are exposed to a wide variety of environmental signals, either inhibitory or stimulatory. Depending on the result of the integration of these signals, the resting state (qNSC) is maintained or the transition to an activated state (aNSC) occurs. Once activated, DG NSCs carry out a series of asymmetric divisions that produce neuron precursors until they eventually differentiate into astrocytes (Encinas et al., 2011); the ratio of NSC to neurons varies in the DG with age, and a trend toward astroglia differentiation can be observed in aged mice resulting in depletion of the NSC pool and reduced neurogenesis (Akers et al., 2014). Thus, the number of hippocampal NSCs decreases with age and at the same time, these cells undergo a transition to a senescent-like state characterized by a complex morphology. The ability of these senescent cells to undergo activation is considerably reduced, remaining inactive for a longer time in the DG of older adults (Diaz-Moreno et al., 2018; Martin-Suarez, Valero, Muro-Garcia, & Encinas, 2019). Recent reports show that gene expression of NSC changes in the aged DG to reduce NSC activation and ensure the maintenance of the stem cells population (Harris et al., 2021) and a pool of long term quiescent stem cells are found in the young brain that is preserved for long periods of time (Ibrayeva et al., 2021). Therefore, the aged niche negatively regulates qNSC activation to preserve the NSC pool reducing neurogenesis. Thus, the search for strategies that promote neurogenesis without depleting the NSC pool could be of use to develop treatments that alleviate the cognitive deterioration associated with aging.

In a previous work, we showed that a diterpene with 12-deoxyphorbol structure (12-desoxyphorbol 13-isobutyrate: DPB or ER272), stimulated hippocampal neurogenesis in adult healthy mice (Dominguez-Garcia et al., 2020), suggesting its ability to act as a pharmacological drug that promotes neurogenesis in mouse models with a compromised cognitive capacity. The mouse model with accelerated aging senescence accelerated mouse prone SAMP8 and its control senescence-accelerated mouse resistant SAMR1 (Takeda, 2009) have long been used to study brain aging and its effects on cognitive ability and neurogenesis. In addition to multisystemic aging, this model presents precociously and progressively cognitive deterioration (Yagi, Katoh, Akiguchi, & Takeda, 1988) starting at 4 months and after 6 months it begins to develop typical characteristics of Alzheimer’s disease (Butterfield & Poon, 2005; Dobarro, Orejana, Aguirre, & Ramirez, 2013; Pallas et al., 2008). Interestingly, in this model, an activation of neurogenesis can be observed at early stages, resulting in the subsequent depletion of the NSC reservoir caused by extracellular signals present in the niche (Diaz-Moreno et al., 2013; Gang et al., 2011; Soriano-Canton et al., 2015). We show in here that chronic treatment of SAMP8 mice with ER272 for two months previous to the initiation of cognitive deterioration improves cognitive performance concomitantly increasing adult hippocampal neurogenesis. The analysis of the cellular and molecular mechanisms of action shows that the compound facilitates the activation of NSC and facilitates the release of the ligand of the epidermal growth factor receptor (EGFR), the transforming growth factor alpha (TGFα) in a mechanism dependent on classical protein kinase C alpha (PKCα).

## Methodology

### Reagents

Isolation and purification of the 12-deoxyphorbol 13-isobutyrate also referred to as DPB (Ezzanad et al., 2021) or ER272 (Dominguez-Garcia et al., 2021; Geribaldi-Doldan et al., 2015) (CAS:25090-74-8) was performed in our laboratory as described in supplementary methods of Geribaldi-Doldan et al. (Geribaldi-Doldan et al., 2015). PKC inhibitors were purchased from Sigma-Aldrich (St. Louis, MO) and Calbiochem (Millipore, Billerica, MA). SmartPool One target siRNA specific for each PKC was obtained from Horizon (Cambridge, UK).

### Animal subjects

Four-month-old SAMR1 and SAMP8 male mice were housed under controlled conditions of temperature (21-23°C) and light (LD 12:12) with free access to food (AO4 standard maintenance diet, SAFE, Épinay-sur-Orge, France) and water. Care and handling of animals were performed according to the Guidelines of the European Union Council (2010/63/EU), and the Spanish regulations (65/2012 and RD53/2013) for the use of laboratory animals. All studies involving animals are reported in accordance with the ARRIVE guidelines for reporting experiments involving animals (Kilkenny et al., 2010; McGrath, Drummond, McLachlan, Kilkenny, & Wainwright, 2010).

### Intranasal administration of ER272

ER272 was delivered intranasally while the animal was placed in a standing position with an extended neck as previously described (Marks, Tucker, Cavallin, Mast, & Fadool, 2009). 18 μL of each solution (1 μM ER272 in saline, or saline as vehicle) was delivered over both nasal cavities alternating 3 μL/each using a micropipette. Mouse was maintained in such position for 10 additional seconds to ensure all fluid was inhaled. In all experiments, mice were coded, treatment (vehicle or ER272) was assigned randomly to code numbers and applied. In addition, blind quantifications were performed to avoid subjective biases.

### Experimental design to test the effect of ER272 on SAMP8 mice

Experimental design is represented in the diagram included in figure 1A. Four-month old SAMR1 and SAMP8 mice were treated with either vehicle or ER272 by intranasal administrations during 8 weeks. SAMR1 mice were used as the control group as indicated in previous reports (Takeda, 2009). During the treatment period, mice received intraperitoneal injections of 5-bromo-2-deoxyuridine (BrdU) every other day. On week 6 mice were subjected to behavioral tests (Morris water Maze, MWM; new object discrimination, NOD and open field) being sacrificed at the end of the 8^th^ week.

**Figure 1.**
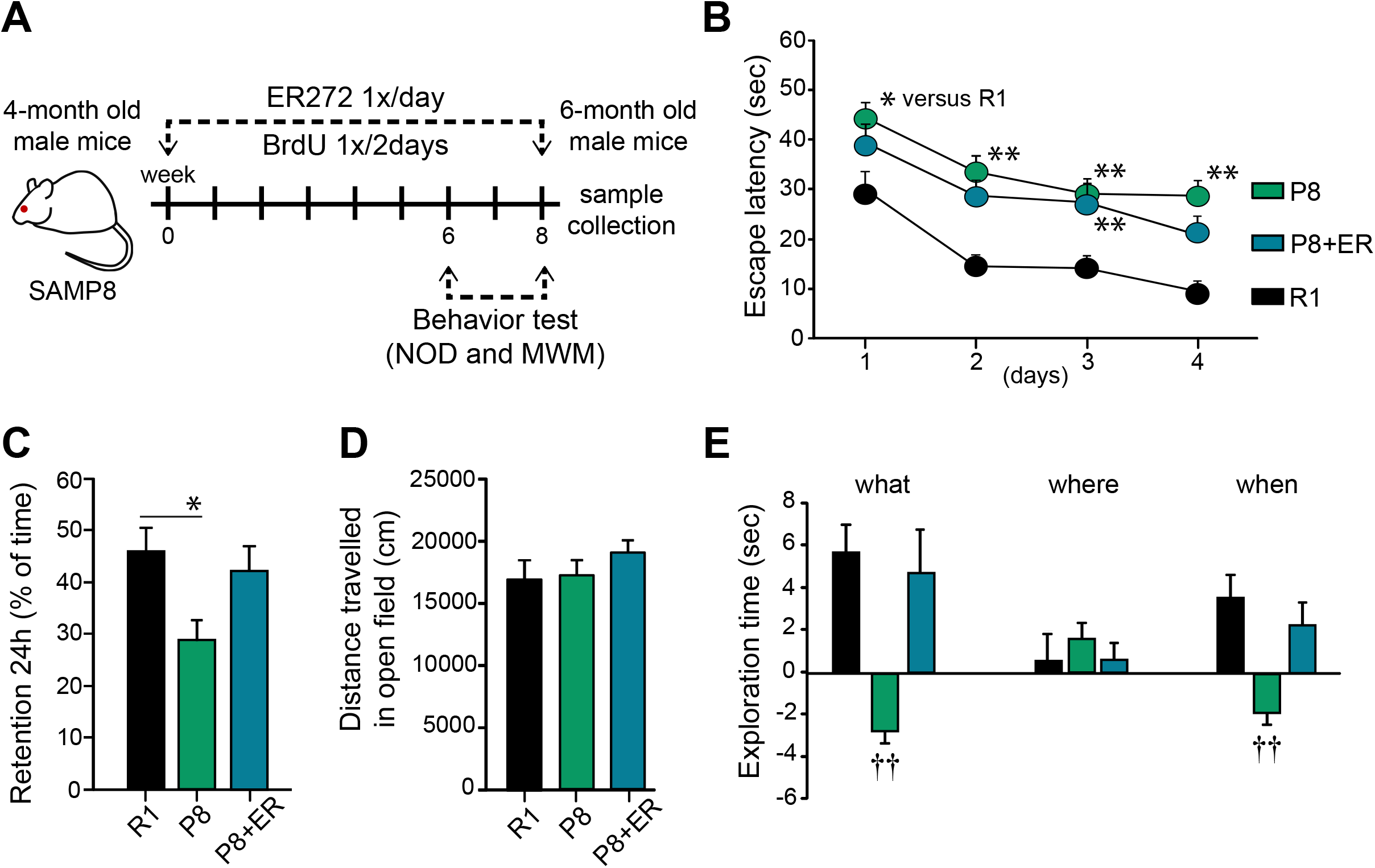
Treatment of SAMP8 mice with ER272 improves cognitive performance. **A)** Experimental design. Four month-old SAMP8 mice (P8) were treated with vehicle or ER272 (ER) for 8 weeks until the age of 6 months. They were compared with SAMR1 mice (R1) of the same age treated with vehicle. Mice received BrdU during the 8 week period every two days. Behavior tests took place during the last two weeks. **B)** Scape latency (sec) in the MWM test. A compromise was observed in SAMP8 male mice when they were evaluated in the MWM and a slight improvement was observed for treated mice (SAMP8-ER). Individual daily assessment revealed a better performance as training sessions progressed: day 1 [F_(2,116)_=4.64, *p=0.031vs. R1], day 2 [F_(2,116)_=10.26, **p<0.01 vs. R1], day 3 [F_(2,113)_=3.67, **p<0.001 vs. R1], day 4 [F_(2,116)_=11.14, **p<0.001 vs. R1]. **C)** Retention test in the MWM. In the 24 h retention phase SAMP8 male mice spent shorter times in the quadrant where the platform used to be located (quadrant 2) [F_(2,25)_=4.16, *p=0.028 vs. R1]. The treatment reverted this situation. **D)** Distance travel in the open field test. No differences were observed among groups when distances travelled in the open field were analyzed [F_(3,38)_=1.83, p=0.158]. **E)** NOD test. Episodic memory was severely affected in male SAMP8 mice when “what” and “when” paradigms were assessed, while ER treatment counterbalanced this situation. “What” [F_(2,84)_=9.55, ††p<0.001 vs. rest of the groups], “where” [F_(2,84)_=0.309, p=0.735], “when” [F_(2,89)_=8.69, ††p<0.001 vs. rest of the groups]. Data are the means ± S.E.M of ten animals, n = 10. Differences detected by one-way ANOVA followed by Tukey b test.

### Motor activity and new object discrimination (NOD) task

Behavioral testing commenced 10 days before sacrifice. Motor activity was analysed measuring the distance travelled by each mouse during a 30 minutes period before initiating the NOD test. One day after the object habituation training (a red-cylinder and a yellow-trapezoid), the NOD test begins. Then integrated episodic memory for the paradigms “what”, “when” and “where” was analyzed as described in previous reports (Ramos-Rodriguez et al., 2013). In brief, next day animals were placed in the centre of the box for 5 minutes with 2 objects (red cilinder and yellow trapezoid) for habituation purposes. The next day mice were placed in the bock with 4 copies of a novel object (blue balls) arranged in a triangle-shaped spatial configuration and allowed to explore them for 5 min. After a delay of 30 min, the mice received a second sample trial with 4 novel objects (red cones), arranged in a quadratic-shaped spatial configuration, for 5 min. After a delay of 30 min, the mice received a test trial with 2 copies of the object from sample trial 2 (“recent” objects) placed at the same position, and two copies of the object from sample trial 1 (“familiar” objects) placed one of them at the same position (“old non-displaced” object) and the another in a new position (“familiar displaced” object). Integrated episodic memory for “what”, “where” and “when” was analyzed as previously described (Dere, Huston, & De Souza Silva, 2005; Infante-Garcia et al., 2018; Segado-Arenas et al., 2018): “What” was defined as the difference in time exploring familiar and recent objects, “where” was defined as the difference in time exploring displaced and non-displaced objects and “when” was defined as the difference between time exploring familiar non displaced and recent non displaced objects.

### Morris water maze (MWM)

Spatial memory and learning tasks were analyzed before sacrifice using the MWM test in control and treated mice the 48 h after the NOD test was concluded as previously described (Ramos-Rodriguez et al., 2013).

### Cerebrospinal fluid extraction

Cerebrospinal fluid (CSF) collection was performed as described by Lim et al., 2018 (Lim et al., 2018). Mice were anesthetized as described above and placed prone on the stereotaxic instrument. Muscles were moved to the side and dura mater over the cisterna magna was exposed. The capillary tube was placed and inserted into the *cisterna magna* through the *dura mater*, lateral to the *arteria dorsalis spinalis*. Finally, the CSF was collected.

### Concentration of TGFα in CSF

TGFα was measured in the CSF using commercial ELISA kits, MBS2508394 (MyBioSourse, Inc, San Diego, CA), following the manufacturer’s instructions. CSF was centrifugated for 20 min at 10g and 4° C; then supernatant was collected. Blanks (diluent only) were included in each independent determination. Blanks were subtracted from measurements before comparisons were made.

### Brain processing and immunohistochemistry

At the end of the treatment brains were perfused with paraformaldehyde (PFA) and-sliced using a cryotome into 30 μm sections. Immunohistochemistry was performed as previously described (Garcia-Bernal et al., 2018; Geribaldi-Doldan et al., 2015; Murillo-Carretero et al., 2017). See antibodies in supplementary table 1.

### Quantification of neurogenesis in brain sections

Cells positive for BrdU, DCX, NeuN, GFAP, SOX2, S100β, in the DG were estimated as described (Rabaneda et al., 2008; Rabaneda et al., 2016). After perfusion, mouse brains were codded and blind quantification was performed as previously described (Geribaldi-Doldan et al., 2018). Positive cells were counted throughout the entire lateral or laterodorsal walls of the lateral ventricles in every fifth section containing the hippocampus; 14-16 sections were analyzed per brain under confocal microscopy (Zeiss LSM 900 Airyscan 2). Confocal imaging was taken every 1.50 μm in the Z-plane using 20X objective. Cell density was calculated for each section relative to the DG volume (mm^3^) and averaged for each animal as reported previously (Dominguez-Garcia et al., 2021).

### Morphological analysis

DCX^+^ cells 3D reconstructions from confocal stack images were obtained using Neurolucida 360 (MBF Bioscience, VT, USA) and the total dendritic length, total dendritic surface, total dendritic segments, number of terminal segments and 3D-Sholl analysis were analyzed with Neurolucida Explorer (MBF Bioscience, VT, USA) according to previous reports (Carrascal, Nieto-Gonzalez, Torres, & Nunez-Abades, 2009; Nunez-Abades, He, Barrionuevo, & Cameron, 1994).

### Western blot

Tissue was homogenized using a commercial lysis buffer (Cell Signaling, USA) supplemented with a protease and phosphatase inhibitor cocktail (Sigma, USA). The homogenates were sonicated and centrifuged at 4° C for 5 min at 16,000 g. Supernatants were collected and protein concentration was measured using the Pierce BCA Protein Assay Kit (Thermo Scientific, Waltham, MA, USA). Equal amounts (30 μg) of total protein from each cellular extract were subjected to SDS-PAGE and western blotting. Proteins were separated on gradient 4-15 % precast polyacrylamide gels (Mini-PROTEAN TGX Stain-Free Protein Gels, BioRad, USA), followed by electrophoretic transfer to PVDF membranes (Schleicher & Schuell, Keene, Netherlands). Membranes were then soaked in blocking buffer (Invitrogen, Carlsbad, CA, USA) for 30 min and incubated overnight at 4° C with primary antibodies (see antibodies in supplementary table 1). Membranes were washed and signal was detected using commercial kits (Western Breeze, Invitrogen, Carlsbad CA, USA) containing either anti-rabbit or anti-mouse secondary antibodies conjugated to alkaline phosphatase, plus the corresponding chemiluminescent substrate following manufacturer’s instructions. Membranes were developed using Chemidoc Touch Imaging System 732BR1030 (BioRad, USA).

### HEK293T culture, cloning and transfection

HEK293T obtained from ATCC (Manassas, VA, USA) were cultured and transfected as previously described (Geribaldi-Doldan et al., 2018). After an overnight incubation, cells were left for 30 min in serum-free Fluorobrite DMEM (Thermo Fisher Scientific, Waltham, MA, USA) and used either in time-lapse experiments

### Cloning of human Neuregulin and TGFα cDNA fused to eGFP and mCherry

Full-length cDNA encoding the membrane-bound isoform of human pro-neuregulin-1 β1-type (NRG1, NCBI reference sequence: NP_039250.2) with mCherry cDNA inserted between nucleotides 93 and 94 of NRG1 open reading frame was cloned into pEGFP-N1 to add EGFP cDNA to the 3’ end. Construct was synthesized by GeneCust (Boynes, France) to generate the mCherry-NRG1-eGFP construct. The mCherry-TGFα-eGFP construct containing the human transforming growth factor alpha (TGFA, NCBI reference sequence: NM_003236.4), containing mCherry cDNA between nucleotides 126 and 127 of TGFA was built using the same strategy and synthesized by GeneCust (Boynes, France).

### siRNA transfection

For the specific inhibition of classical PKCα and β cells were transfected with specific siRNA Smart Pool siRNA One target (Horizon, Cambridge, UK). Cell transfection with TGFα construct and siRNA pool were performed 18 hours after seeding; for this, cells were changed to antibiotic-free medium and transfected using Lipofectamine 2000 (Invitrogen, Carlsbad, CAD, USA), following the manufacturer’s instructions. Lipofectamine was removed 6 hours later.

### Time lapse experiments and fluorescence analysis of recombinant mCherry-TGFα-eGFP protein in the culture medium of HEK293T

HEK293T cells were plated in μ–dishes (35 mm high, Ibidi, Munich, Germany) and treated with ER272 and/or inhibitors, and images were taken as described in figure legends.

### Statistical analysis

The data and statistical analysis comply with the recommendations on experimental design and analysis in pharmacology (Curtis & Abernethy, 2015). Statistical analysis was performed using the computer program IBM SPSS Statistics 22. Unless otherwise indicated, normal distribution of the data was first analyzed using a Shapiro-Wilks test. Then, a Brown Forsythe test was performed to test the equality of variances. Afterwards, when more than one treatment group were compared, statistical analyses were performed using one-way ANOVA followed by a post-hoc Tukey’s test unless otherwise indicated. Two way-ANOVA (group x day) was used in the acquisition phase of the MWM. Differences were considered significant at values of p<0.05. In general, sample size used in statistical analysis were n = 6-10 for *in vivo* experiments, n = 5-9 for *in vitro* experiments and n = 9-12 in behavioral tests. Sample sizes were chosen based on previous works related to this one (Carrasco et al., 2014; Dominguez-Garcia et al., 2021; Geribaldi-Doldan et al., 2018; Murillo-Carretero et al., 2017; Rabaneda et al., 2016).

## RESULTS

### Long-term intranasal administration of ER272 improves cognitive performance in SAMP8 mice

As described previously, spatial memory was compromised in male SAMP8 mice as reflected in the evaluation on the MWM and a slight improvement was observed in SAMP8 mice treated with ER272. No significant group x day effect was detected [F_(2,462)_=0.358, p=0.905], however individual daily assessment revealed a better performance of treated mice as training sessions progressed (Fig. 1 B). In the retention phase of the MWM, we observed that SAMP8 mice spent significantly shorter times in the quadrant where the platform used to be located when compared with SAMR1 mice. Interestingly, in the retention phase SAMP8 mice treated with ER272 did not show differences in comparison with the SAMR1 control group (Fig. 1 C). As expected, episodic memory was severely affected in male SAMP8 mice when “what” and “when” paradigms were assessed (p<0.001), while ER272 treatment counterbalanced this situation. On the contrary, no differences were observed in the “where” paradigm (Fig. 1 E). As control, no differences were observed in the distance travelled in the open field (Fig. 1 D).

### Long term intranasal administration of ER272 promotes neurogenesis in the DG of SAMP8 mice

Mice were given intranasal administrations of ER272 for eight weeks and received BrdU injections every two days starting on the first day of treatment (Fig. 1 A). The analysis of BrdU^+^ cells did not show a reduction in the total number of cells that had incorporated BrdU when SAMP8 mice were compared with SAMR1 (Fig. 2 A-B, F). However, the DG of SAMP8 treated mice showed a 2-fold increase in the number of cells that have incorporated BrdU over the course of the treatment compared to SAMP8 mice (Fig. 2 A-C, F). In the control, most of the BrdU^+^ cells were DCX^+^ (Fig. 2 A-B F, G). These percentages were lower in the SAMP8 mice but not if the SAMP8 mice were treated with ER272 (Fig. 2 A-C, F, G). The analysis of total DCX^+^ cells in all groups revealed that at six months of age, SAMP8 mice showed a lower number of DCX^+^ cells (Fig. 2 A-B, E) compared to SAMR1. Interestingly, the treatment of SAMP8 mice with ER272 increased the number of DCX^+^ cells by 2-fold compared to SAMP8 and compensated this reduction (Fig 2 B-C, E). Identically, the number of DCX^+^ cells that had incorporated BrdU over the course of two months was dramatically reduced in SAMP8 mice compared to control (Fig. 2 A-B, G), whereas this number increased by 2-fold in mice treated with ER272 (Fig. 2 B-C, G).

**Figure 2.**
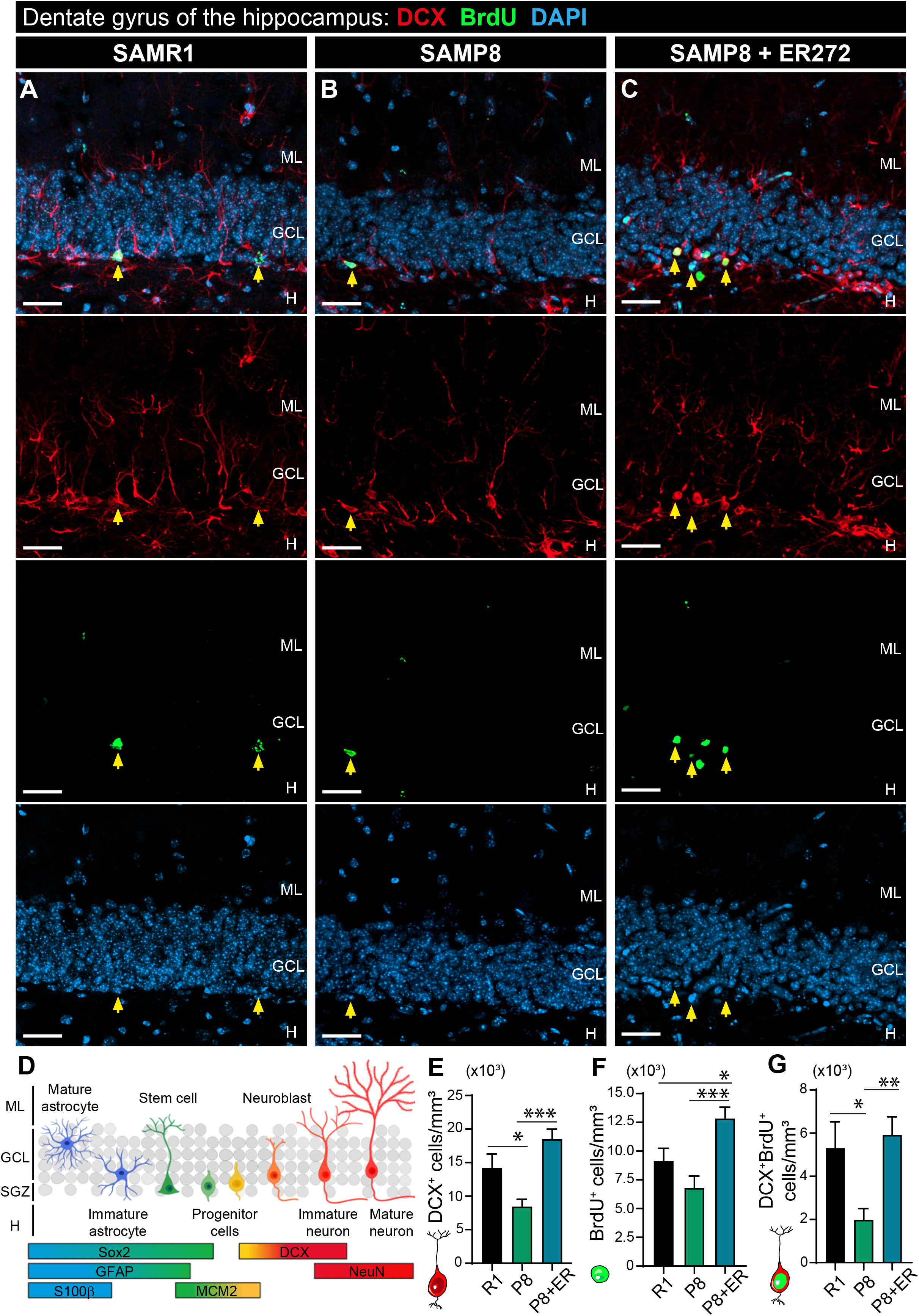
Effect of long-term intranasal administration of ER272 to SAMP8 mice on DCX^+^ neuroblasts. **A-C)** Representative confocal microcopy images of the DG of the hippocampus of six-month-old SAMR1 (R1) and SAMP8 (P8) and male mice treated with vehicle (A, B respectively) or SAMP8 mice treated with ER272 (P8+ER) (C) during eight weeks as indicated in Fig. 1 A. Slices were processed for the immunohistochemical detection of the proliferation marker BrdU (lower medium panel; green) and DCX (upper medium panel; red). DAPI staining is shown in blue (lower panel). Merged channels are shown in the upper panel. Yellow arrows indicate DCX^+^BrdU^+^DAPI^+^ cells. **D)** Schematic drawing showing the hierarchical model of adult hippocampal neurogenesis indicating the different markers along the cell lineage within the dentate gyrus. **E)** Graph shows the total number of DCX^+^ in the DG of the hippocampus per mm^3^ [F_(2,29)_= 10.88, *p=0.036 R1vs P8] [F_(2,29)_= 10.88, *p<0.001 P8 vs P8+ER]. **F)** Graph shows the total number of BrdU^+^ nuclei in the DG of the hippocampus per mm^3^ [F_(2,32)_=10.58, *p=0.045 R1 vs P8+ER] [F_(2,32)_= 10.58, ***p<0.001 P8 vs P8+ER]. **G)** Graph shows the total number of BrdU^+^DCX^+^ cells in the DG of the hippocampus per mm^3^ [F_(2,31)_=6.898, *p=0.034 R1 vs P8] [F_(2,31)_=6.898, **p<0.003 P8 vs P8+ER]. Data are the means± S.E.M of six animals, n =6. Differences detected by one-way ANOVA followed by Tukey b test. Scale bar represents 25 μm.

The number of NeuN^+^ cells was not statistically different between control and SAMP8 group at six months of age, however, a 2-fold increase in the number of NeuN^+^ cells was observed in mice treated with ER272 (Fig. 3 A-D). In addition, a 2-fold increase in the number of NeuN^+^ cells that had incorporated BrdU was observed (Fig. 3 A-C, E). In control mice, the number of DCX^+^ cells that expressed NeuN was reduced compared to SAMR1, but this number was increased by 2-fold in SAMP8 mice treated with ER272 (supplementary figure S1).

**Figure 3.**
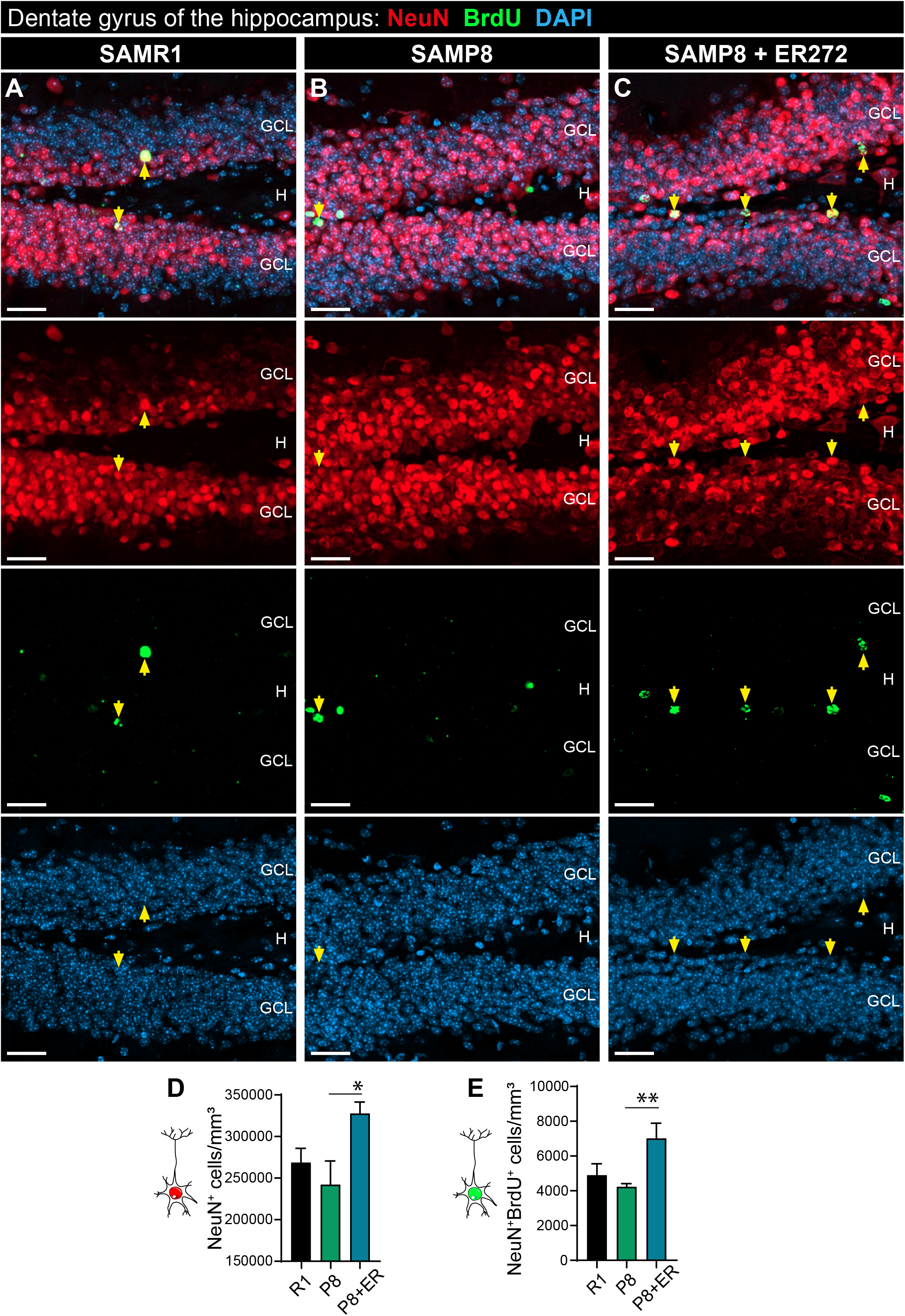
Effect of long-term intranasal administration of ER272 to SAMP8 mice on NeuN^+^ cells. **A-C)** Representative confocal microcopy images of the DG of the hippocampus of six-month-old SAMR1 (R1) and SAMP8 (P8) and male mice treated with vehicle (A, B respectively) or SAMP8 mice treated with ER272 (P8+ER) (C) during eight weeks as indicated in Fig. 1 A. Slices were processed for the immunohistochemical detection of the proliferation marker BrdU (lower medium panel; green) and NeuN (upper medium panel; red). DAPI staining is shown in blue (lower panel). Merged channels are shown in the upper panel. Yellow arrows indicate NeuN^+^BrdU^+^DAPI^+^ cells. **D)** Graph shows the number of NeuN^+^ nuclei in the DG of the hippocampus per mm^3^ [F_(2,14)_=3.934, *p=0.038 P8 vs P8+ER]. **E)** Graph shows the number of NeuN+BrdU+ nuclei in the DG of the hippocampus per mm^3^ [F_(2,8)_=11.04, **p=0.004 P8 vs P8+ER]. Data are the means±S.E.M of six animals, n =6. Differences detected by one-way ANOVA followed by Tukey b test. Scale bar represents 25 μm.

### Morphology of DCX^+^ cells is altered in SAMP8 mice and restored by ER272 treatment

The study of the morphological properties of DCX^+^ cells revealed that the total dendritic length, the total dendritic surface, the total dendritic segments and the number of terminal segments were reduced in SAMP8 mice compared to control (Fig. 4 A-C, D-G). Also, the number of dendritic segments on the 4ry and 5ry centrifugal order were reduced in SAMP8 mice compared to SAMR1 (Fig. 4 A-C, H, I). It was very interesting to note that ER272 treatment reverted all these age-induced morphological alterations (Fig. 4 A-I). In addition, Sholl analysis shows a lower number of intersections at distances from soma of 60-70 μm in SAMP8 mice compared to control (Fig. 4, J), an effect that was reverted by the ER272 treatment.

**Figure 4.**
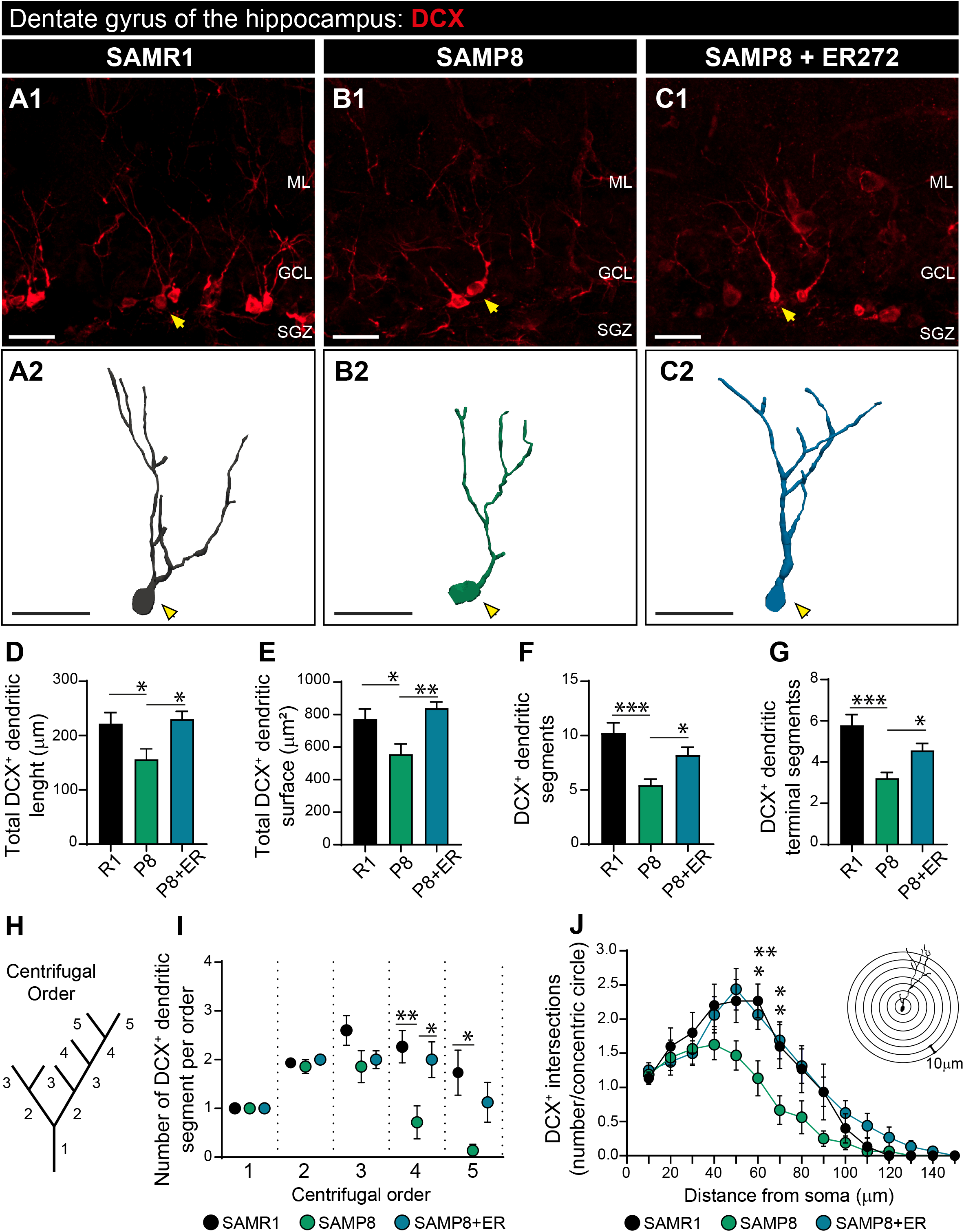
Altered morphology of DCX^+^ cells in SAMP8 is restored by ER272 treatment. **A-C)** Representative confocal microcopy images of the DG of the hippocampus of six-month-old SAMR1 (R1) and SAMP8 (P8) and male mice treated with vehicle (A1, B1 respectively) or SAMP8 mice treated with ER272 (P8+ER) (C1) during eight weeks. Examples of the reconstructed DCX^+^ cells used in the analysis of SAMR1 and SAMP8 treated with vehicle (A2, C2 respectively) and SAMP8 treated with ER272 (C2). Yellow arrows indicate DCX^+^ cells used as 3D representative reconstruction. **D-G)** Total dendritic length [F_(2,43)_=4.94, *p=0.040 R1 vs P8] [F_(2,43)_=4.94, *p=0.016 P8 vs P8+ER] (D), total dendritic surface [F_(2,43)_=7.157, *p=0.024 R1 vs P8] [F_(2,43)_=7.157, **p=0.002 P8 vs P8+ER] (E), total dendritic segments [F_(2,40)_=9.51, ***p<0.001 R1 vs P8] [F_(2,40)_=9.51, *p=0.038 P8 vs P8+ER] (F) and the number of terminal segments [F_(2,41)_=10.5, ***p<0.001 R1 vs P8] [F_(2,41)_=10.5, *p=0.045 P8 vs P8+ER] (G) were measured using Neurolucida Explorer. **H)** Scheme representing the centrifugal order of different dendritic segments. **I)** Number of DCX^+^ segments in the different centrifugal orders. Quaternary order [F_(2,42)_=5.50, **p<0.009 R1 vs P8] [F_(2,42)_=5.50, *p=0.032 P8 vs P8+ER]. Quinary order [F_(2,43)_=4.76, *p<0.011 R1 vs P8]. **J)** Sholl analysis showing the number of intersections of the DCX^+^ dendritic segments at different distances from soma of cells. 60 μm from soma [F_(2,43)_=6.292, **p<0.005 R1 vs P8] [F_(2,43)_=6.292, *p<0.022 P8 vs P8+ER]. 70 μm from soma [F_(2,42)_=4.917, *p<0.026 R1 vs P8] [F_(2,42)_=4.917, *p<0.025 P8 vs P8+ER]. A total of 15 cells were analyzed per group. Data are the means±S.E.M of 15 cells per group, n = 15. Differences detected by one-way ANOVA followed by Tukey b test. Scale bar represents 25 μm.

### Effect of ER272 treatment on niche astrocytes

We next investigated whether astrogliogenesis in these senescent mice was altered and the effect of our drug on the generation of new astrocytes. Results show a larger number of astrocytes GFAP^+^S100β^+^ cells in SAMP8 mice compared to SAMR1 (Fig. 5 A-E). Interestingly, the treatment with ER272 reverted this phenotype. As shown in figure 5 A-E, the treatment of SAMP8 mice with ER272 reduced by 2-fold the number of astrocytes generated during the two-month period.

**Figure 5.**
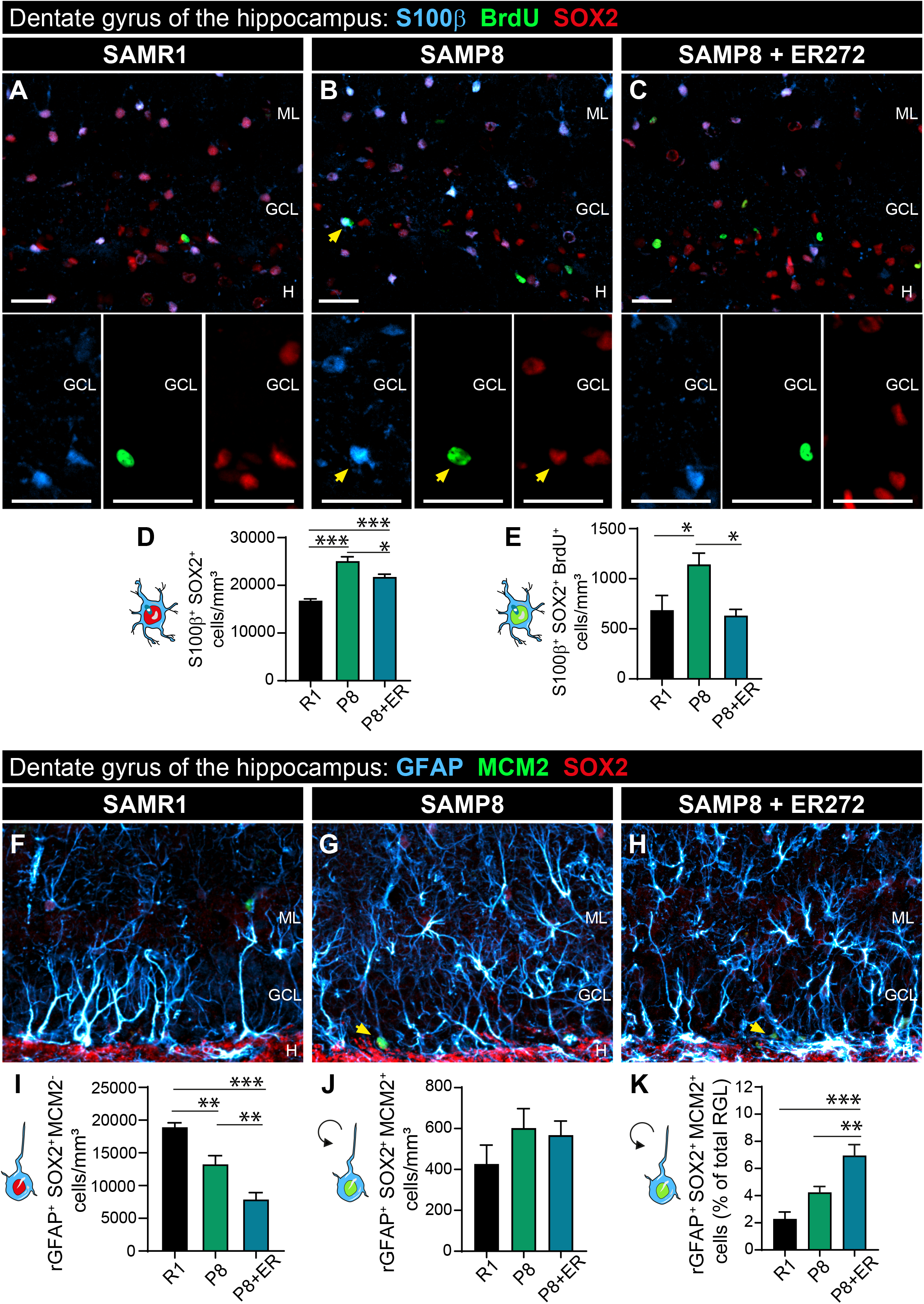
Intranasal administration of ER272 decreases the number of newly generated astrocytes in the dentate gyrus of SAMP8 mice and promotes the radial glial-like cells activation in the dentate gyrus of SAMP8 mice. **A-C)** Representative confocal microcopy images of the DG of the hippocampus of six-month-old SAMR1 (R1) and SAMP8 (P8) and male mice treated with vehicle (A, B respectively) or SAMP8 mice treated with ER272 (P8+ER) (C) during eight weeks as indicated in Fig. 1 A. Slices were processed for the immunohistochemical detection of the proliferation marker BrdU (green), SOX2, a transcription factor expressed in astrocytes (red) and the marker for astrocytes S100β (blue). Yellow arrows indicate S100β^+^BrdU^+^SOX2^+^ cells. **D)** Graph shows the number of S100β+SOX+ cells in the DG per mm^3^ [F_(2,13)_=46.0, ***p<0.001 R1 vs P8] [F_(2,13)_=46.0, ***p<0.001 R1 vs P8+ER] [F_(2,13)_=46.0, *p=0.013 P8 vs P8+ER]. **E)** Graph shows the number of S100β+SOX2+BrdU+ cells in the DG of the hippocampus per mm^3^ [F_(2,13)_=5.48, *p<0.036 R1 vs P8] [F_(2,13)_=5.48, *p=0.036 P8 vs P8+ER]. **F-H)** Representative confocal microcopy images of the DG of the hippocampus of six-month-old SAMR1 and SAMP8 and male mice treated with vehicle (F, G respectively) or SAMP8 mice treated with ER272 (H) during eight weeks as indicated in Fig. 1 A. Slices were processed for the immunohistochemical detection of the Glial Fibrillary Acidic Protein GFAP (blue), SOX2 (red) and the cell cycle marker MCM2 (green). Yellow arrows indicate rGFAP^+^MCM2^+^SOX2^+^ cells. **I)** Graph shows the number of rGFAP^+^SOX2^+^MCM2^-^ with a radial glial-like cell (r) morphology in the DG per mm^3^ [F_(2,15)_=30.5, **p<0.003 R1 vs P8] [F_(2,15)_=30.5, ***p<0.001 R1 vs P8+ER] [F_(2,15)_=30.5, **p=0.005 P8 vs P8+ER]. **J)** Graph shows the number of rGFAP^+^SOX2^+^MCM2^+^ radial glial-like cells in the DG of the hippocampus per mm^3^. **K)** Graph shows the percentage of rGFAP^+^SOX2^+^MCM2^+^ radial glial-like cells per rGFAP^+^SOX2^+^ in the DG of the hippocampus [F_(2,15)_=14.3, ***p<0.001 R1 vs P8+ER] [F_(2,15)_=14.3, *p=0.019 P8 vs P8+ER]. Data are the means± S.E.M of six animals, n =6. Differences detected by one-way ANOVA followed by Tukey b test. Scale bar represents 25 μm.

### Effect of ER272 treatment on neural stem cells (radial glial like cells, rNSC)

We next investigated whether the treatment affected the NSC and its activation. Particularly we analyzed the number of cells that expressed the glial marker protein glial fibrillary acidic protein (GFAP) and the transcription factor SOX2 (GFAP^+^SOX2^+^) that showed a radial glial like morphology (rGFAP^+^SOX2^+^). We could observe that the total number of rGFAP^+^SOX2^+^ cells decreased in SAMP8 mice compared with SAMR1 mice. Also, the treatment of SAMP8 mice with ER272 reduced this number (Fig. 5 F-I). To analyze the proportion of NSC undergoing cell cycle, we studied the number of rGFAP^+^SOX2^+^ that expressed the mitosis marker MCM2 (rGFAP^+^SOX2^+^MCM2^+^). We did not observe a higher number of rGFAP^+^SOX2^+^MCM2^+^ cells in SAMP8 compared to control on the 6^th^ month of age neither the proportion of rGFAP^+^SOX2^+^MCM2^+^ cells was different in SAMP8 mice compared to SAMR1 (Fig. 5 F-H, J) and although the treatment with ER272 did no change the number of rGFAP^+^SOX2^+^MCM2^+^ cells in SAMP8 treated with ER272 in comparison with SAMP8 it significantly increased the proportion of rGFAP^+^SOX2^+^MCM2^+^ of the total rGFAP^+^SOX2^+^ (Fig. 5 F-H, K).

### ER272 promotes TGFα release in a reaction dependent on classical PKCα

Previous reports had indicated that activation of protein kinase C facilitated the ADAM17-mediated release of TGFα *in vitro*. To understand the mechanism of action of ER272, we studied its capacity to facilitate the release of the growth factor TGFα using time lapse imaging. As shown in figure 6, pro-TGFα was cloned in a mammalian expression vector flanked by red fluorescent protein in the N-terminal portion and eGFP in the C-terminal (Fig. 6 A-B). We transfected cultures of HEK293T cells with this construct and using time lapse imaging we quantified the release of the TGFα ligand from the pro-ligand in the presence or absence of ER272 and in cultures co-transfected with siRNA to block the expression of different classical PKC isozymes. The release of TGFα results in the loss of red fluorescence and therefore, in a reduction in the red/green fluorescence ratio (Fig. 2 B). As shown in figure 6 C-G, the addition of ER272 to cultures expressing the double fluorescent protein labeled construct, dramatically reduced the ratio red/green over the course of three hours (Fig 6. C, G and supplementary movie) whereas no changes were observed when the vehicle was added to the culture (Fig. 6 C, D and supplementary movie). Addition of ER272 to cells in which PKCα expression had been abolished did not show any change in the fluorescent ratio (Fig. 6 C, E and supplementary movie) and the addition of ER272 to cells in which PKCβ expression had been inhibited showed an intermediate effect on the red/green fluorescence ratio (Fig. 6 C, F and supplementary movie).

**Figure 6.**
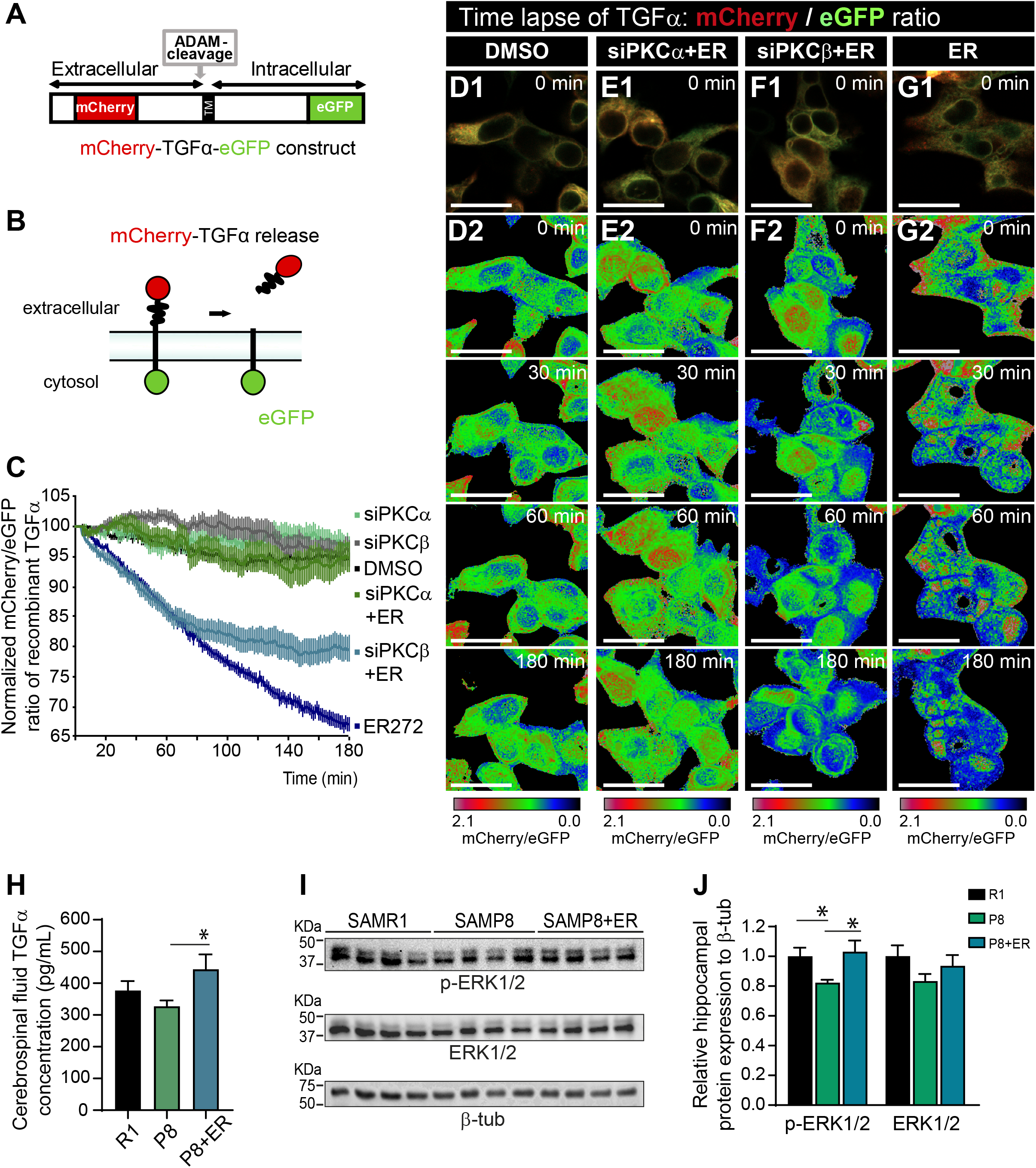
Molecular mechanisms underlying the effects of ER272: role of TGFα. A) Scheme of mCherry-TGFα-eGFP construct B) Mechanisms of TGFα-bound fluorescence release. C) Quantitative analysis of the microscopic images obtained from the time-lapse assays of HEK293T cells expressing mCherry-TGFα-eGFP and stimulated with ER272, PKCα siRNA, and/or PKCβ siRNA. mCherry/eGFP ratios were normalized to the average mCherry/eGFP ratio measured before stimulation. The mean normalized mCherry/eGFP ratios and SEM are shown, n = 40. See also supplementary movie 1. D-G) mCherry/eGFP ratio images at the indicated time points are shown in the intensity modulated display-mode. The color ranging goes from red to blue to represent mCherry/eGFP ratios. The upper and lower limits of the ratio range are shown on the right. Scale bar represents 25 μm. H) Concentration of TGFα in the cerebrospinal fluid (CSF) measured by ELISA [F_(2,9)_=4.09, *p=0.045 P8 vs P8+ER]. I) Images of chemiluminescence signal of immunoblot detection of pERK1/2, total ERK1/2, and β-tubulin (loading control) in the hippocampus of SAMR1 (R1), SAMP8 (P8) treated with vehicle and SAMP8 treated with ER272 (P8+ER). J) Integrated optical density quantification of images obtained from immunoblots (see supplementary figure S2): hippocampal protein expression of pERK1/2 [F_(2,17)_=6.59, *p=0.029 R1 vs P8] [F_(2,17)_=6.59, *p=0.010 P8 vs P8+ER] and total ERK1/2 according to β-tubulin. Data in J are the means± S.E.M of eight animals, n =8. Differences detected by one-way ANOVA followed by Tukey b test.

### Effect of ER272 on intracerebral concentration of TGFα and TGFα initiated signaling events

Thus, we next investigated the effect of ER272 on the release of TGFα *in vivo*. To do that, we extracted cerebrospinal fluid (CSF) from the *cisterna magna* of SAMR1, SAMP8 and SAMP8 treated mice before sacrifice and analyzed the concentration of TGFα in the CSF by ELISA immunodetection (Fig. 6 H). We observed that the concentration of TGFα in the CSF of mice treated with ER272 was significantly higher than in control and SAMP8 groups (Fig. 6 H). Also, the analysis by western blot of the activation of the Map kinase pathway, by looking at ERK1/2 phosphorylation, revealed that the phosphorylation of ERK1/2 is reduced in SAMP8 mice compared to SAMR1 mice and the treatment with ER272 reverts this effect (Fig. 6 I-J and supplementary figure S2).

## DISCUSSION

Increasing age is linked to cognitive decline and it is an independent risk factor for the development of neurodegenerative disorders (Murman, 2015). Specific types of memory have been found affected in aged individuals (Gray & Barnes, 2015), episodic memory being the most affected in the aging population (Nyberg et al., 2012). The search for therapeutic strategies that prevent cognitive decline is nowadays a matter of crucial significance. In this context, strategies targeting the positive regulation of adult hippocampal neurogenesis may result in the development of new treatments to delay cognitive decline in the elderly. We had previously reported the isolation of molecules that promote neurogenesis in the adult brain and suggested its use as pharmacological tools to treat cognitive deficiencies related to a reduction in neurogenesis such as the diterpene with 12-deoxyphorbol structure ER272 (Geribaldi-Doldan et al., 2015; Murillo-Carretero et al., 2017). This molecule facilitate the secretion of trophic factors *in vitro* that act on receptor tyrosine kinases of the ErbB1-4 family (Dominguez-Garcia et al., 2020).

To analyze the effect of ER272 on a murine model of pathological aging we have used the senescence model SAMP8 using as a control the SAMR1 strain (Takeda, 2009). SAMP8 recapitulate the transition from healthy aging to Alzheimer’s disease since these mice spontaneously develops memory and learning deficiencies (Takeda, 2009) exhibiting Alzheimer’s disease like features: oxidative stress, increased amyloid precursor protein (APP) and its mRNA as well as and amyloid ß (Aß) levels, phosphorylation of tau, and astrogliosis among others) starting at six months of age (Butterfield & Poon, 2005; Diaz-Moreno et al., 2013; Morley et al., 2000; Pallas et al., 2008). Studies on the neurogenesis in SAMP8 mice reveal that they display a proliferative response that precedes the AD phenotype (Diaz-Moreno et al., 2013) and an accelerated depletion of the hippocampal NSC pool compared with SAMR1 that coincides in space and time with an increase in astroglial differentiation and a reduction in neurogenesis (Diaz-Moreno et al., 2018).

Using this model, we show that as previously described, SAMP8 mice show cognitive impairment (Takeda, 2009), particularly, we show that spatial memory is compromised showing difficulties to perform in the MWM test. Similarly, episodic memory in these mice is also impaired as it can be inferred from their performance in the NOD test. These results agree with previous results, which show similar memory impairments in this model (Dobarro et al., 2013). Interestingly, treatment of SAMP8 mice with ER272 reverts the NOD impairment and improves their capacity to perform in the MWM test, being part of the novelty of this work the identification of a compound that is able to revert spatial and episodic memory impairment.

The study of hippocampal neurogenesis in these mice shows that the generation of new hippocampal neurons is also compromised in this model of pathological aging. Previous results show a significant increase in proliferation in the DG of two-month-old SAMP8 animals compared to SAMR1 that returns to control levels in older mice (Diaz-Moreno et al., 2013). Accordingly, we do not see any changes in the proliferation of cells (BrdU^+^) in the DG of six-month-old SAMP8 mice compared to SAMR1, however, we observe a robust increase in proliferation in SAMP8 mice that have been treated with ER272 from the 4^th^ to the 6^th^ month of age, indicating that the proliferative response initiated before two months of age is maintained in treated mice as a consequence of the treatment. Interestingly, on the 6^th^ month we observe a reduction in the number of neuroblasts (DCX^+^ cells) and in the number of DCX^+^ neuroblasts that incorporated BrdU over the course of the 2 months before sacrifice in SAMP8 mice compared to SAMR1. The treatment of mice with ER272 avoided this reduction indicating the capacity of this compound to potentiate the generation of neuroblasts.

It was quite noticeable the aberrant morphology of the DCX^+^ cells on the SAMP8 mice compared to the SAMR1 control group. These results complement previous ones, which show an abnormal accumulation of the DCX^+^ cell population in two-month-old SAMP8 mice. The location of DCX^+^ cells was altered, these cells migrated deeper into the granule cell layer and were aberrantly positioned (Diaz-Moreno et al., 2018). We show in here an aberrant morphology of DCX^+^ cells in the GCL (granular cell layer) characterized by shorter dendrites, a lower number of dendritic segments and a general retraction of the dendritic tree in six-month-old SAMP8 mice. However, the treatment of mice with ER272 prevents the development of these aberrant morphological alterations found in the aged mouse brain.

The reduction in the number of DCX^+^ neuroblasts in SAMP8 mice was not concomitant with a reduction in the activation of NSC (GFAP^+^SOX2^+^MCM2^+^ cells). In this context, an explanation for the lower number of DCX^+^ cells could be that we also detected an increase in the number of newly generated astrocytes S100β^+^BrdU^+^ suggesting that not all of the activated NSC are producing neuroblasts but they are generating astrocytes. Previous reports have shown that astroglial differentiation of neural stem cells is a hallmark of neurogenesis in the aged brain also indicating that NSC are lost in aged mice due to their preferential differentiation into mature astrocytic cells (Diaz-Moreno et al., 2018; Encinas et al., 2011). We observe in here that indeed an elevated number of mature astrocytes can be observed in six-month-old SAMP8 mice compared to SAMR1 and it is noticeable the increase in the number of astrocytes that have incorporated BrdU within the eight-week period prior to sacrifice. Notwithstanding, the treatment of SAMP8 mice for eight weeks prevented this phenomenon reducing the number of BrdU^+^ astrocytes to that found in SAMR1 mice and increasing the number of DCX^+^ neuroblasts.

Noteworthy was the fact that we despite the reduction in DCX^+^ cells in the SAMP8 mouse compared to SAMR1 no differences were found in the number of NeuN^+^ neurons or NeuN^+^ neurons that incorporated BrdU over the course of the 2 months before sacrifice. These puzzling results agree with the previous reports that show in five-month-old SAMP8 that the number of NeuN^+^ cells that have incorporated BrdU (administered daily) over the last month of life is not different from that in SAMR1 mice (Sasaki et al., 2020). An explanation for the lack of differences in the number of neurons that have been generated in the SAMP8 is that survival rate of newly generated neurons in the SAMP8 mice might be higher than in SAMR1 or that differentiation is accelerated as a response. Interestingly, the number of DCX^+^ cells that expressed NeuN was also reduced in SAMP8 mice compared to SAMR1, indicating that a higher rate of differentiation may not be the reason for the lack of differences in NeuN^+^BrdU^+^ cells and suggesting that in SAMP8 the survival rate of NeuN^+^ cells may be higher than in SAMR1. The treatment of SAMP8 mice with ER272 led to a higher number of DCX^+^ and NeuN^+^ cells that had incorporated BrdU as well as a higher number of DCX^+^/NeuN^+^ cells life indicating that this pharmacological agent was positively regulating neurogenesis by facilitating differentiation and probably not affecting survival.

Previous studies show that the proportion of NSC decreases with age in the mouse brain reducing their senescent cells ability to undergo activation and remaining inactive or quiescent for prolonged periods of time (Diaz-Moreno et al., 2018; Harris et al., 2021; Ibrayeva et al., 2021; Martin-Suarez et al., 2019), thus preserving the NSC reservoir from exhaustion. However, for the continuous generation of new neurons a certain basal firing rate is required (Ziebell, Dehler, Martin-Villalba, & Marciniak-Czochra, 2018). We have found in here that in the DG of SAMP8 mice a reduction in the number of NSC is observed with age, as demonstrated by the lower number of radial glial like cells that express SOX2 and GFAP. This reduction is concomitant with an increase in astrogliosis. A similar depletion of NSC has previously been observed in the DG of six-month-old SAMP8 mice compared to SAMR1 mice accompanied with a response in NSC activation (Diaz-Moreno et al., 2018). Using our experimental model this activation is slightly higher but this increase does not reach statistisc significance. It was clear, though that the treatment of mice with ER272 increased the number of activated NSC compared to non-treated SAMP8 mice and with SAMR1, thus indicating that this treatment activates NSC and reduces the NSC pool. However, although the maintenance of quiescence is probably a determining factor when it comes to preserving the neurogenic rate during aging because it protects the NSC reservoir from total exhaustion, the generation of new neurons that we observed in treated mice improves cognitive performance supporting that for this continuos generation of neurons a certain basal activation rate is required (Ziebell et al., 2018).

Finally, in order to understand the mechanisms of action that lead to the effect of ER272, we have analyzed its role on the release of TGFα to the extracellular medium. TGFα is one of the physiological ligands of the EGFR and as other EGFR ligands it is expressed as a membrane bound pro-ligand with the EGFR binding domain in the extracellular N-terminal portion of the protein. The shedding of the EGFR binding domain that releases the ligand is catalyzed by a metalloprotease of the ADAM family, ADAM17, via the regulation of protein kinase C (Dang et al., 2013; Dang et al., 2011). Using a fusion protein construct in which pro-TGFα was cloned in frame with a eGFP protein in the C-terminal end and a mCherry protein in the N-terminal, we show that in culture cells, ER272 facilitates the release of TGFα as shown in previous reports (Dominguez-Garcia et al., 2020). This release is impaired in cells in which the expression of classical protein kinase C alpha (PKCα) has been abolished, indicating that ER272 stimulated the PKCα dependent release of TGFα. Accordingly, in SAMP8 mice treated with ER272 for two months, we observe an increase in the concentration of TGFα in the CSF. This increase is concomitant with the stimulation of the MAPK/ERK1/2 signaling cascade. The activation of this pathway is reduced in SAMP8 mice and the treatment with ER272 restores this activity. This finding agrees with a previous report in which SAMP8 mice show a reduced pERK1/2 activity that is related to changes in cognitive performance and neuronal morphology (Vasilopoulou et al., 2022). This suggests that an increase in TGFα may be one of the mechanisms involved in the positive regulation of neurogenesis found in SAMP8 mice avoiding cognitive impairment. The implication of TGFα in cognitive performance has previously been suggested agreeing with our findings (Alipanahzadeh et al., 2014; Vasilopoulou et al., 2022).

In summary, several pieces of evidence show the existence of alterations in neurogenesis in aged mice (Diaz-Moreno et al., 2018; Kempermann, Kuhn, & Gage, 1998; van Praag, Shubert, Zhao, & Gage, 2005), some of them suggesting that neurogenesis does not occur in the neuropathologically aged DG as it does in the non-aged niche. As previously reported, we have observed that neurogenesis (Diaz-Moreno et al., 2018) and cognitive performance (Dobarro et al., 2013; Takeda, 2009) is impaired in the brain of a mice model of pathological aging. Nevertheless, we report the effect of the diterpene with 12-deoxyphorbol structure ER272, a pharmacological compound that positively regulates adult hippocampal neurogenesis by increasing the proportion of activated NSC, favoring the generation of neuroblasts, neuronal differentiation and reorganizing neurogenesis. Thus, our results suggest that this compound helps rejuvenating the DG niche. Concomitantly, we show an effect of this drug improving cognitive performance in the same model. The study of the mechanism of action reveals that an increase in TGFα release via the activation of PKCα may be the underlying mechanism of action. It could then be concluded that we have found a small molecule that may work as a pharmacological drug to positively regulate neurogenesis and cognitive performance in the pathologically aged brain.

## Acknowledgments

We thank the Servicio de experimentation y producción animal (SEPA) de la Universidad de Cádiz as well as the Servicios Centrales de apoyo a la investigación en Ciencias de la Salud (SCICS) and Servicios centrales de Ciencia y tecnología (SC-ICYT) de la Universidad de Cádiz.

## Conflict of Interest

The authors declare that the research was conducted in the absence of any commercial or financial relationships that could be construed as a potential conflict of interest.

## Funding

This work was supported by the Spanish Ministerio de Ciencia, Innovatión y Universidades (grant number RTI-2018-099908-B-C21 and RTI-2018-099908-B-C22 granted to CC and RH-G), by the Consejería de Economía, Conocimiento, Empresas y Universidades (grant number FEDER-UCA18-106647 granted to CC) and by the Consejería de Salud y Familias 80% co-finaced by EDRF ITI regional funds (ITI-Cadiz-0042-2019)

## Author Contribution

**RG-O, PN-A, MG-A:** experimental design, data acquisition and analysis, discussion of results article preparation and writing.

**SM-O, IA-N, CB, SD-G, NG-D, CV:** data acquisition and analysis, discussion of results,

**RH-G and AE:** isolation characterization and preparation of chemical compound

**CC:** conception of the work, experimental design, data acquisition and analysis, discussion of results, article preparation and writing, funding acquisition.

## Data availability statement

Data will be available to interested person upon request

## Notes

### Competing Interest Statement

The authors have declared no competing interest.

