## supplementary figures and tables for "Rescue of neurogenesis and age-associated cognitive decline in SAMP8 mouse: role of transforming growth factor alpha"

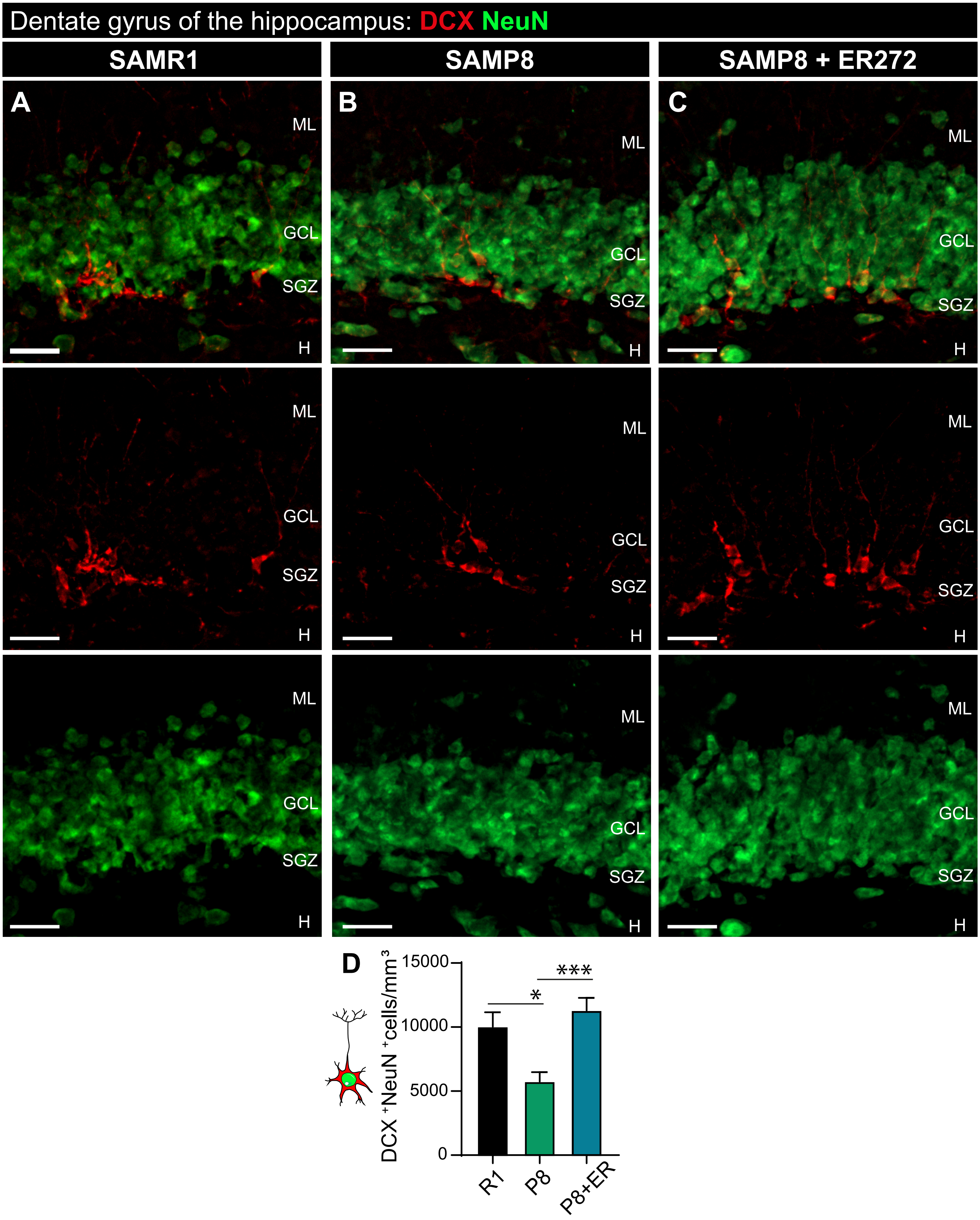


**Supplementary Figure S1. Effect of long-term intranasal administration of ER272 to SAMP8 mice on DCX+NeuN+ cells. A-C)** Representative confocal microcopy images of the DG of the hippocampus of six-month-old SAMR1 (R1) and SAMP8 (P8) and male mice treated with vehicle (A, B respectively) or SAMP8 mice treated with ER272 (P8+ER) (C) during eight weeks as indicated in Fig. 1 A. Slices were processed for the immunohistochemical detection of DCX (medium panel; red) and NeuN (lower panel; green) markers. **D)** Graph shows the number of DCX+NeuN+ cells in the DG of the hippocampus per $mm^3$ [$F(2,30)=8.977$, *$p<0.011$ R1 vs P8] [$F(2,30)=8.977$, ***$p<0.001$ P8 vs P8+ER]. Data are the means ± S.E.M of six animals, n = 6. Differences detected by one-way ANOVA followed by Tukey b test. Scale bar represents 25 µm.

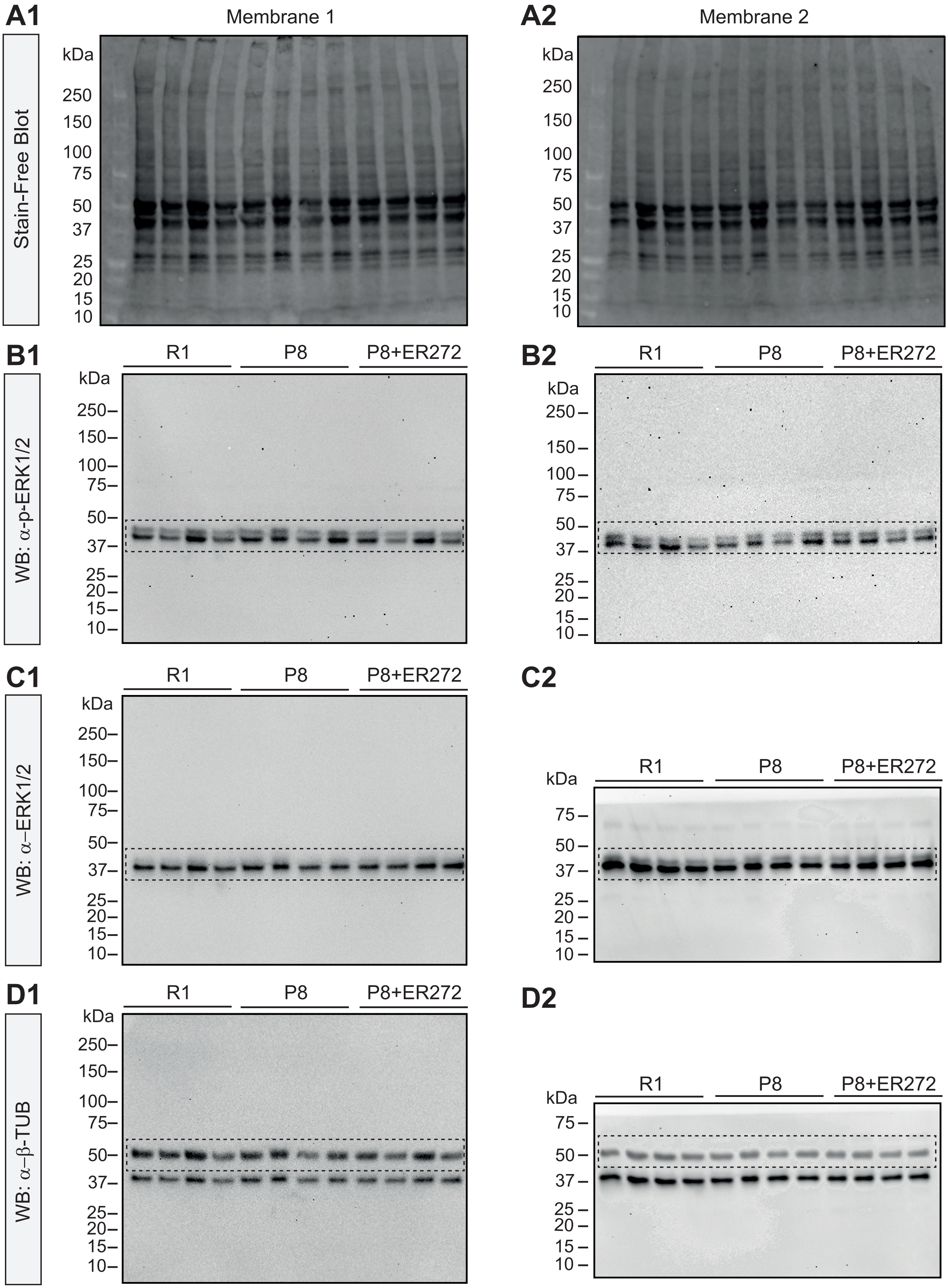


**Supplementary Figure S2. Total hippocampal protein membrane imaging and antibody detection. A)** Total hippocampal proteins of six-month SAMR (R1) and SAMP8 (P8) male mice treated with vehicule or SAMP8 mice treated with ER272 (P8+ER), separated on 4-15% polyacrylamide gels and transferred to PVDF membranes (membrane 1 and membrane 2). **B)** Images of chemiluminescence signal of immunoblot detection of p-ERK1/2 (dotted squares). B2 image conrresponds to representative blots in the main figure. **C)** Images of chemiluminescence signal of immunoblot detection of total ERK1/2 (dotted squares). C2 image conrresponds to representative blots in the main figure. **D)** Images of chemiluminescence signal of immunoblot detection of loading control β-tubulin (dotted squares). D2 image conrresponds to representative blots in the main figure. n=8.

| Antibody | Host | Isotype | Epitope retrieval | Staining pattern | Source | Reference |
|---|---|---|---|---|---|---|
| Anti-DCX | Rabbit | Polyclonal | DCX, neuroblast marker | Cytoplasmic | Abcam (Cambridge, UK) | ab18723 |
| Anti-S100β | Rabbit | Polyclonal | S100β, astrocyte marker | Cytoplasmic | Abcam (Cambridge, UK) | ab41548 |
| Anti-NeuN | Rabbit | Monoclonal | NeuN, neuronal marker | Nuclear | Abcam (Cambridge, UK) | ab177487 |
| Anti-phospho-p44/42 MAPK (ERK1/2) | Rabbit | Polyclonal | Phospho-p44/42 MAPK (ERK1/2), marker of stimulation of MAPK (ERK1/2) signaling cascade | Cytoplasmic/ nuclear | Cell Signaling (Danvers, MA, USA) | 9101 |
| Anti-SOX2 | Rabbit | Polyclonal | SOX2, astrocyte/stem cell marker | Nuclear | Santa Cruz Biotechnology, Santa Cruz, CA, USA) | sc-20088 |
| Anti-SOX2 | Goat | Polyclonal | | Nuclear | RD Systems Minneapolis, CO, USA) | AF2018 |
| Anti-MCM2 | Mouse | Monoclonal | MCM2, cell proliferation marker | Nuclear | BD Bioscience ( East Rutherford, NJ, USA) | 610701 |
| Anti-p44/42 MAPK (ERK1/2) | Mouse | Monoclonal | p44/42 MAPK (ERK1/2), marker of total MAPK (ERK1/2) | Cytoplasmic/ nuclear | Cell Signaling (Danvers, MA, USA) | 4696 |
| Anti- β-tubulin | Mouse | Monoclonal | β-tubulin, core protein in microtubules, loading control for Western Blot | Cytoplasmic | Thermo Fisher Scientific (Waltham, MA, USA) | MA5-16308-HRP |
| Anti-GFAP | Chicken | Polyclonal | GFAP, glial marker | Cytoplasmic | Abcam (Cambridge, UK) | ab4674 |
| Anti-NeuN | Rat | Monoclonal | NeuN, neuronal marker | Nuclear | Abcam (Cambridge, UK) | ab279297 |
| Anti-BrdU | Rat | Monoclonal | BrdU, cell proliferation marker | Nuclear | Abcam (Cambridge, UK) | ab6362 |

**Primary antibodies supplementary table 1:** List of primary antibodies used in the study. Specifying host, isotype, epitope retrieval, staining pattern, source and reference.

| Antibody | Host | Dilution | Fluorescence | Source | Reference |
|---|---|---|---|---|---|
| Alexa Flour anti-rabbit | Donkey | 1:1000 | 647 | Invitrogen (Carlsbad, CA, USA) | A-32795 |
| Alexa Flour anti-rabbit | Donkey | 1:1000 | 594 | Invitrogen (Carlsbad, CA, USA) | A-21207 |
| Alexa Flour anti-rabbit | Donkey | 1:1000 | 488 | Invitrogen (Carlsbad, CA, USA) | A-21206 |
| Alexa Flour anti-goat | Donkey | 1:1000 | 647 | Invitrogen (Carlsbad, CA, USA) | A-11058 |
| Alexa Flour anti-mouse | Donkey | 1:1000 | 488 | Invitrogen (Carlsbad, CA, USA) | A-21206 |
| Alexa Flour anti-chicken | Goat | 1:1000 | 647 | Abcam (Cambridge, UK) | ab150175 |
| Alexa Flour anti-rat | Donkey | 1:1000 | 488 | Life Technologies (Eugene, OR, USA) | A-21208 |

**Secondary antibodies supplementary table 2:** List of secondary antibodies used in the study. Specifying host, dilution used, fluorescence conjugated, source and reference.

| Intercalating agent | Target | Staining pattern | Source | Reference |
|---|---|---|---|---|
| DAPI | Double-stranded DNA binding dye | nuclear | (Sigma, St. Louis, MO, USA) | D9542 |

**DNA staining:** DNA-intercalanting-dye used to stain cell nuclei. Specifying target, staining pattern, source and reference.
